# Maize EHD1 is Required for Kernel Development and Vegetative Growth through Regulating Auxin Homeostasis

**DOI:** 10.1101/390831

**Authors:** Yafei Wang, Wenwen Liu, Hongqiu Wang, Qingguo Du, Zhiyuan Fu, Wen-Xue Li, Jihua Tang

**Affiliations:** National Key Laboratory of Wheat and Maize Crop Science, Collaborative Innovation Center of Henan Grain Crops, College of Agronomy, Henan Agricultural University, Zhengzhou, 450002, China; National Engineering Laboratory for Crop Molecular Breeding, Institute of Crop Science, Chinese Academy of Agricultural Sciences, Beijing 100081, China

## Abstract

The roles of EHDs in clathrin-mediated endocytosis (CME) in plants are poorly understood. Here, we isolated a maize mutant, designated as *ehd1*, which showed defects in kernel development and vegetative growth. Positional cloning and transgenic analysis revealed that *ehd1* encodes an EHD protein. Internalization of the endocytic tracer FM4-64 was significantly reduced in *ehd1* mutant and *ZmEHD1* knock-out mutants. We further demonstrated that ZmEHD1 and ZmAP2 σ subunit physically interact in the plasma membranes. Cellular IAA levels were significantly lower in *ehd1* mutant than in wild-type maize. Auxin distribution and ZmPIN1a-YFP localization were altered in *ehd1* mutant. Exogenous application of 1-NAA but not GA3 rescued the seed germination and seedling emergency phenotypic defects of *ehd1* mutants. Taken together, these results indicate that ZmEHD1 regulates auxin homeostasis by mediating CME through its interaction with the ZmAP2 σ subunit, which is crucial for kernel development and vegetative growth of maize.

## Introduction

Endocytosis, which can be clathrin-dependent or clathrin-independent, is the internalization of plasma membrane (PM) proteins or uptake of extracellular materials for transport to the endosome (Murphy et al., 2005). Clathrin-mediated endocytosis (CME) is the major gateway into plant cells (Fan et al., 2015). Although detailed information is available on the roles of endocytosis in animals, researchers have only recently documented that CME is critical for various developmental processes in plants, including cell polarity determination (Van et al., 2011; Wang et al., 2013), male reproductive organ development (Blackbourn and Jackson, 1996; Kim et al., 2013), gametophyte development (Backues et al., 2010), and embryogenesis (Fan et al., 2013; Kitakura et al., 2011). CME is also important in plant responses to abiotic and biotic stresses by internalizing of PM-resident transporters or receptors, such as the boron transporter (BOR1) (Takano et al., 2010), the iron-regulated transporter (IRT1) (Barberon et al., 2011), the auxin efflux transporter PIN-FORMED (PINs) (Dhonukshe et al., 2007), the brassinosteroid receptor BRASSINOSTEROID INSENSITIVE 1 (BRI1) (Di et al., 2013), the bacterial flagellin receptor FLAGELIN SENSING 2 (FLS2) (Robatzek et al., 2006), and the ethylene-inducing xylanase receptor (LeEIX2) (Bar and Avni, 2009).

CME is a complex process that can be divided into at least four steps: the budding of a vesicle, the packaging of cargo into the vesicle, the release of the mature vesicle from the PM, and the fusing of the vesicle with the endosome (Sigismund et al., 2012). Because clathrin cannot directly bind to the PM or to cargoes, the formation of clathrin-coated endocytic vesicles initiates the association of adaptor protein complex 2 (AP2) with the PM, and thus AP2 plays a central role in the initiation of CME. The AP2 complex forms a heterotetrameric complex containing two large subunits (β1-5 and one each of γ/α/δ/ε/ζ), a medium subunit (μ1-5), and a small subunit (σ1-5) (McMahon and Boucrot, 2011). Our understanding of AP2 in plants has been enhanced by the following recent findings: knockdown of the two *Arabidopsis AP2A* genes or overexpression of a dominant-negative version of the medium AP2 subunit, *AP2M*, was shown to significantly impair BRI1 endocytosis and to enhance brassinosteroid signaling (Di et al., 2013); α and μ adaptins were demonstrated to be crucial for pollen production and viability, as well as elongation of staminal filaments and pollen tubes by modulating the amount and polarity of PINs (Kim et al., 2013); σ adaptin was found to play important roles in the assembly of a functional AP2 complex, and *ap2 σ* loss-of-function *Arabidopsis* exhibited defects in multiple aspects of plant growth and development (Fan et al., 2013).

In addition to the classical AP2 complex, other accessory proteins, including the C-terminal Eps15 homology domain (EHD) proteins, also connect the cargo and membrane lipid to form the CME vesicle. The EHD family contains one member in *Drosophila* and *C. elegans* and four orthologs in mouse and human. All members contain an EHD with two calcium-binding helix-loop-helix motifs (EF-hands), a P-loop with a predicted ATP/GTP binding site, a dynamin-N motif with a nucleotide-binding domain, and a coil-coil region (Bar et al., 2008). The involvement of EHD-containing proteins in CME was described in detail in mammalian cells, in which RME1 was demonstrated to be a key player controlling the recycling of internalized receptors from the endocytic recycling compartment to the PM (Lin et al., 2001). By homologous cloning, two *EHD* genes were isolated from *Arabidopsis* (Bar et al., 2008). The two AtEHD proteins regulated endocytosis in distinct patterns. Over-expression of *AtEHD2* had an inhibitory effect on endocytosis, while down-regulation of *AtEHD1* caused retardation of entry of endocytosed material into plant cells (Bar et al., 2008). However, the mechanisms by which EHD family members linked to CME in plants have not been characterized.

Maize ranks first in total production among major staple cereals and is also an important raw material for biofuel and many other industrial products (Mclaren, 2005). As an important model system for basic biological research, maize has contributed significantly to our knowledge of plant development and evolution, and this understanding has been used to elucidate the developmental mechanisms of maize kernel morphogenesis. The earliest genetic studies performed in maize included analyses of defective kernel (*dek*) mutants. Although several hundred *dek* mutants have been isolated, relatively few corresponding genes have been molecularly cloned because embryo-lethal alleles are difficult to study (Scanlon and Takacs, 2009).

To better understand maize endosperm filling and maturation, we characterized a maize mutant *ehd1* that is impaired in kernel development and vegetative growth. Positional cloning of *ehd1* and transgenic analysis identified the causative locus as a gene that encodes an EHD-containing protein and that is herein designated as *ZmEHDl*. We show that ZmEHD1 is involved in CME through interaction with the ZmAP2 σ subunit. We further demonstrate that *ZmEHD1* affects the gene expression involved in auxin-related processes and that exogenous 1-NAA application rescues the fertility of the *ehd1* mutant. Our results indicate that, by regulating auxin homeostasis, ZmEHD1-mediated CME is crucial for kernel development and vegetative growth of maize.

## Results

### The *ehd1* mutant is impaired in kernel development and vegetative growth

The *ehd1* mutant was originally isolated as a shrunken kernel mutant in the screening of natural mutagenesis defective in the filling of maize grains. At 14 days after pollination (DAP), the kernels of *ehd1* homozygous mutant (*ehdl/ehdl*) and the wild-type (WT) (*EHD1/EHD1* and *EHD1/ehd1*) resembled each other (Figure 1A). At 16 DAP, the WT kernels were canary-yellow while the *ehd1* mutants were ivory-white (Figure 1A). At maturity, both the endosperm and embryo of *ehd1* mutant were shrunken (Figure 1B), which led to a significant reduction in the 100-kernel weight (Figure 1C). The 100-kernel weight of the WT (26.9 g) was 3-times greater than that of the *ehd1* mutant (Figure 1C).

**Figure 1.**
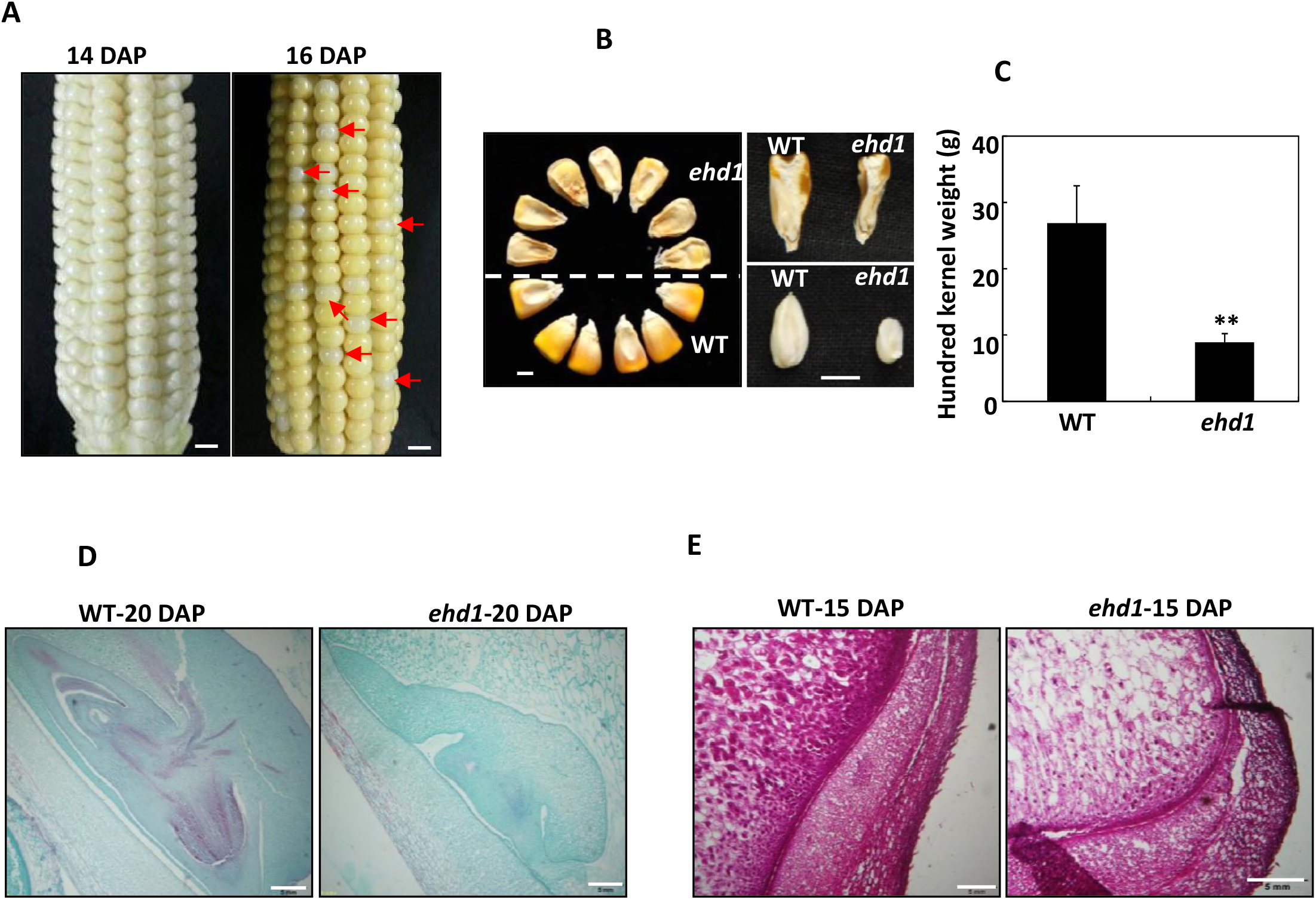
Phenotypes of the maize *ehd1* mutant kernels. (A) F_2_ ears of *ehd1* × Xun9058 at 14 or 16 days after pollination (DAP). The red arrows indicated the *ehd1* kernels. Scale bars = 1 cm. (B) The mature kernels of wild type (WT) and *ehd1* mutant randomly selected from F_2_ ears of *ehd1* × Xun9058. Scale bars = 0.5 cm. (C) 100-grain weight of the WT and the *ehd1* mutant. Values are means and standard errors (n=4). ** indicates a significant difference (P < 0.01) between the WT and the *ehd1* mutant. (D) Comparison of WT and *ehd1* embryos at 20 DAP. The sections stained with Safranin and Fast Green. Scale bars = 5 mm. (E) Microstructure of WT and *ehd1* endosperms at 15 DAP. The sections stained with fuchsin. Scale bars = 5 mm.

To analyze the kernel development of the *ehd1* mutant and the WT, we examined immature embryos at 20 DAP by light microscopy. Longitudinal sections indicated that embryo development was slower in the *ehd1* mutant than in the WT (Figure 1D). Endosperm development and texture were also observed by light microscopy. Compared to WT endosperm cells, those of the *ehd1* mutant had less dense cytoplasm and fewer starch granules (Figure 1E).

### A paper-culture system was used to measure the germination rates of the WT and

the *ehd1* mutant. The germination rate of the *ehd1* mutant was only about 3%, which was far lower than that of the WT (Figure 2A). Relative to the WT seedlings, the germinated *ehd1* seedlings had shorter and fewer primary roots (Figure 2B). Most of the *ehd1* seedlings died before the first two leaves had completely expanded (Figure 2C). These results indicated that *ZmEHD1* is essential for maize kernel development and vegetative growth.

**Figure 2.**
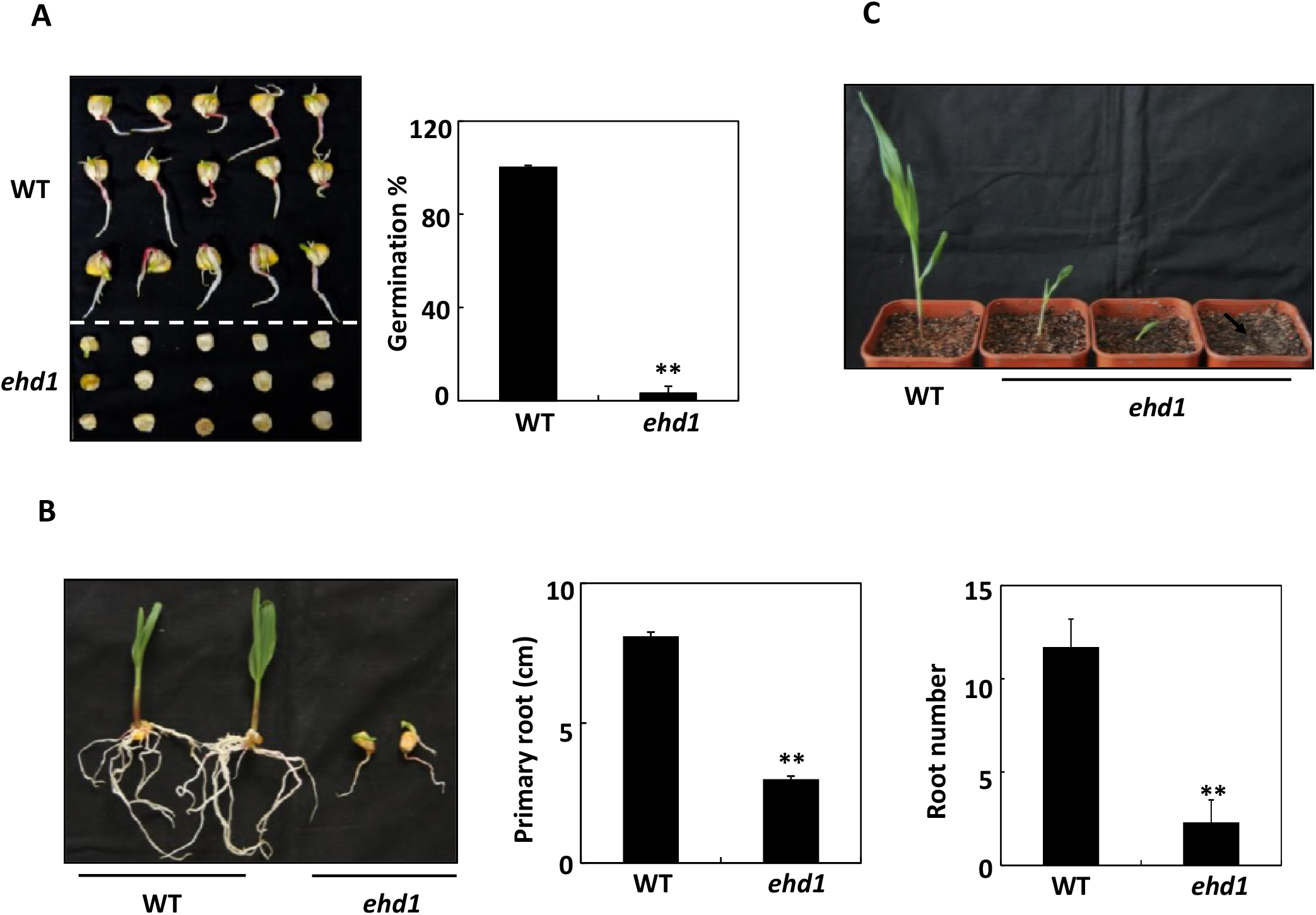
*ZmEHD1* is required for normal growth and development of maize. (A) Germination of the WT and the *ehd1* mutant. Values are means and standard errors of approximately 100 seeds from three independent experiments. (B) Root number and root elongation of the WT and the *ehd1* mutant. Values are means and standard errors ** (n=5). indicates a significant difference (P < 0.01) between the WT and the *ehd1* mutant. (C) Phenotypes of WT and *ehd1* seedlings (n=30). Representative photograph is shown.

The mutant was crossed to an inbred line Xun9058. The hybrid F_1_ displayed normal kernel as Xun9058. Due to the low germination rate of the *ehd1* mutant, the normal kernels from F_2_ ears were planted. Among the 244 F_2_ individuals, 79 had the normal kernel phenotype (*EHD1*/*EHD1*), and 165 had the segregation phenotype (*EHD1/ehd1)*. The segregation ratio followed 1:2 theoretical ratio predicted by a Chi-square test at a 0.05 level (Supplemental Table 1). For each F_2_ individual with kernel phenotype segregation, the pattern of shrunken kernel phenotype to normal kernel phenotype fit a 1:3 segregation ratio (Supplemental Table 2). A F_3_ population with 376 individuals derived from normal kernels of F_2_ selfed ears was also planted to validate the kernel phenotype segregation. Among them, 130 individuals had kernel phenotype (EHD1/EHD1), and 246 had the segregation phenotype (*EHD1/ehd1*) (Supplemental Table 1). These results indicate that this shrunken kernel mutant is controlled by a single recessive gene.

**Table 1.**
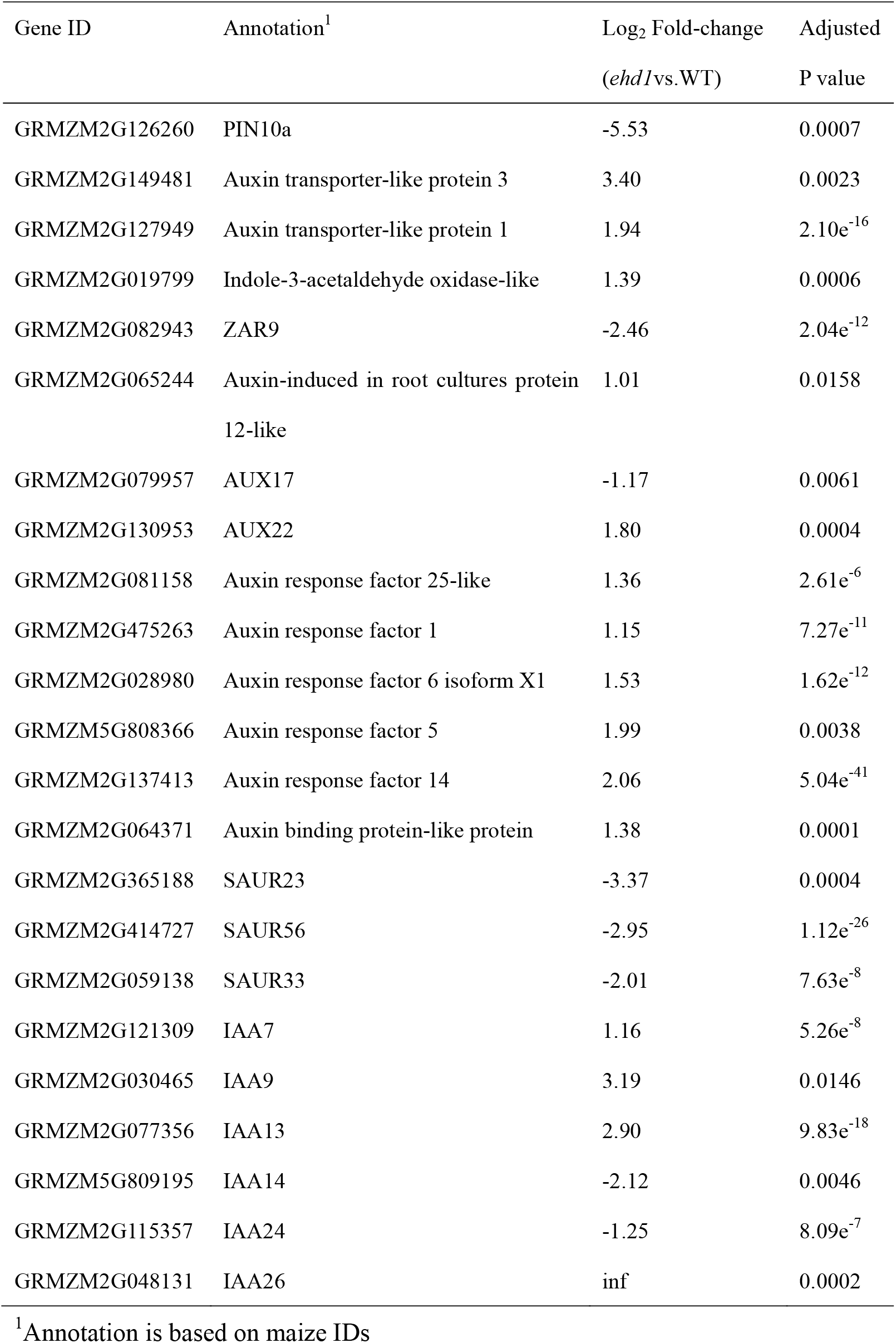
Differentially expressed genes involved in auxin-related processes in the *ehd1* mutant.

| Gene ID | Annotation<sup>1</sup> | Log<sub>2</sub> Fold-change<br>(*ehd1* vs. WT) | Adjusted<br>P value |
| --- | --- | --- | --- |
| GRMZM2G126260 | PIN10a | -5.53 | 0.0007 |
| GRMZM2G149481 | Auxin transporter-like protein 3 | 3.40 | 0.0023 |
| GRMZM2G127949 | Auxin transporter-like protein 1 | 1.94 | 2.10e<sup>-16</sup> |
| GRMZM2G019799 | Indole-3-acetaldehyde oxidase-like | 1.39 | 0.0006 |
| GRMZM2G082943 | ZAR9 | -2.46 | 2.04e<sup>-12</sup> |
| GRMZM2G065244 | Auxin-induced in root cultures protein<br>12-like | 1.01 | 0.0158 |
| GRMZM2G079957 | AUX17 | -1.17 | 0.0061 |
| GRMZM2G130953 | AUX22 | 1.80 | 0.0004 |
| GRMZM2G081158 | Auxin response factor 25-like | 1.36 | 2.61e<sup>-6</sup> |
| GRMZM2G475263 | Auxin response factor 1 | 1.15 | 7.27e<sup>-11</sup> |
| GRMZM2G028980 | Auxin response factor 6 isoform X1 | 1.53 | 1.62e<sup>-12</sup> |
| GRMZM5G808366 | Auxin response factor 5 | 1.99 | 0.0038 |
| GRMZM2G137413 | Auxin response factor 14 | 2.06 | 5.04e<sup>-41</sup> |
| GRMZM2G064371 | Auxin binding protein-like protein | 1.38 | 0.0001 |
| GRMZM2G365188 | SAUR23 | -3.37 | 0.0004 |
| GRMZM2G414727 | SAUR56 | -2.95 | 1.12e<sup>-26</sup> |
| GRMZM2G059138 | SAUR33 | -2.01 | 7.63e<sup>-8</sup> |
| GRMZM2G121309 | IAA7 | 1.16 | 5.26e<sup>-8</sup> |
| GRMZM2G030465 | IAA9 | 3.19 | 0.0146 |
| GRMZM2G077356 | IAA13 | 2.90 | 9.83e<sup>-18</sup> |
| GRMZM5G809195 | IAA14 | -2.12 | 0.0046 |
| GRMZM2G115357 | IAA24 | -1.25 | 8.09e<sup>-7</sup> |
| GRMZM2G048131 | IAA26 | inf | 0.0002 |

<sup>1</sup>Annotation is based on maize IDs

### Positional cloning of *ZmEHD1*

Preliminary genetic mapping of *ZmEHD1* was carried out by BSA using the F_2_ population, which contained 165 individuals that differed in seed size in the ears. The *ZmEHD1* gene was first mapped to a 2,339-kb region between the simple sequence repeat (SSR) marker umc1650 (17 recombinants) and umc1716 (27 recombinants) on the long arm of chromosome 4 (Figure 3A). Because the germination rate of the *ehd1* mutant was rather low, expanded F_3_ mapping population containing ~53,000 normal kernels (*EHD1/EHD1* and *EHD1/ehd1*) was obtained to increase mapping resolution as described by Zuo et al. (2015). Thirteen new polymorphic SSR markers were developed (Supplemental Table 3), and *ZmEHD1* was eventually mapped to a ~6-kb region between the SSR markers RM7 (56 recombinants) and RM13 (5 recombinants) (Figure 3A). This region contains two predicted open reading frames (ORFs) (Figure 3A). Sequence analysis revealed three deletions in the 5’ UTR and eight single-nucleotide polymorphisms (SNPs) in the ORF of GRMZM2G052740 (Figure 3B). Among the eight SNPs in the ORF of GRMZM2G052740, three led to amino acid replacement between the *ehd1* mutant and the WT (Figure 3B). In addition to GRMZM2G052740, GRMZM2G052720 was also located in this region. Its candidacy as *ZmEHD1* gene was excluded because there was no sequence difference between the WT and the *ehd1* mutant. Thus, we inferred that GRMZM2G052740 was the candidate gene for the *ZmEHD1* locus.

**Figure 3.**
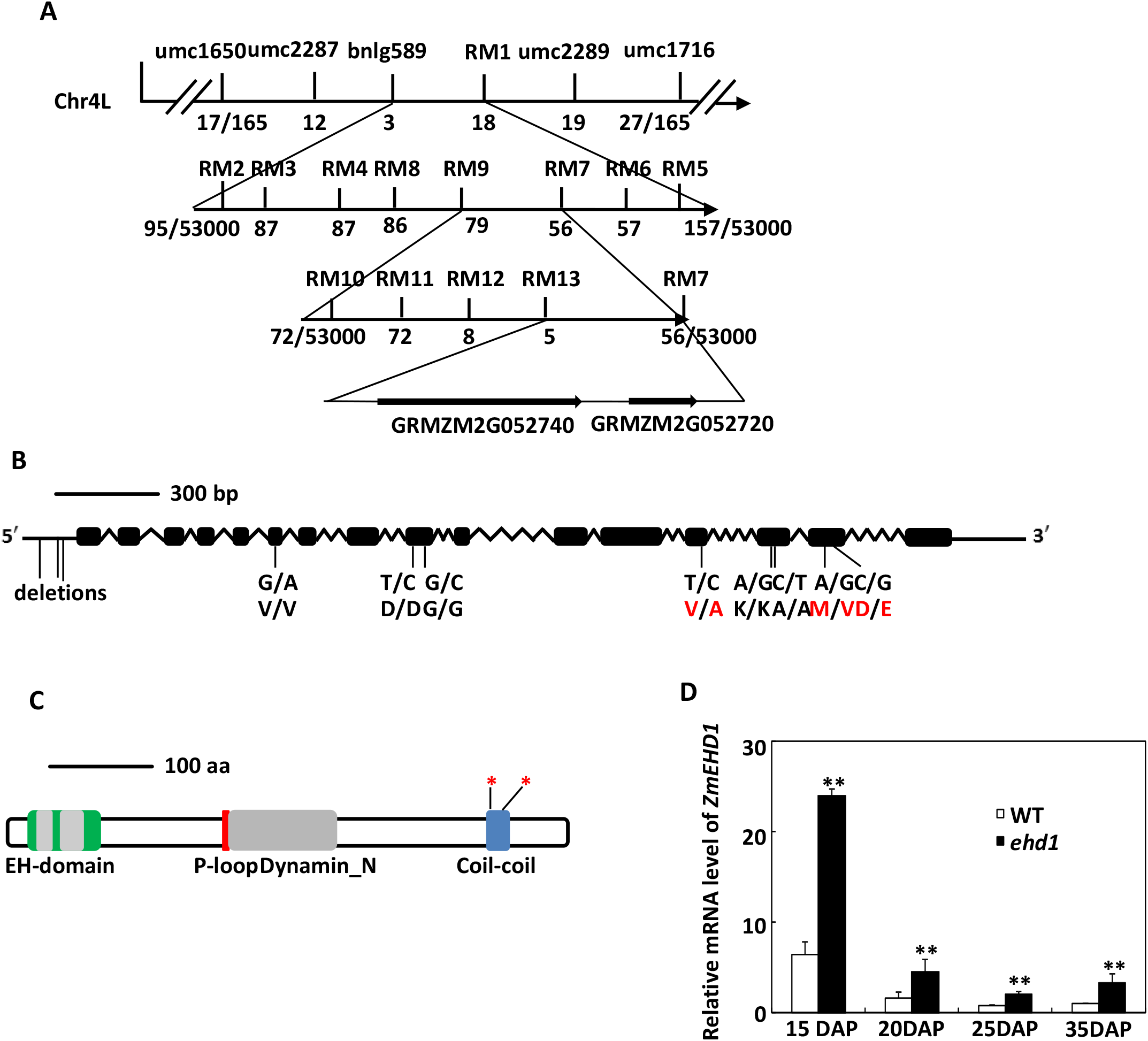
Map-based cloning of *ZmEHD1*. (A) Schematic representation of the positional cloning of *ZmEHD1* gene on chromosome 4. The SSR markers, umc1716 and umc1650, were used for rough mapping. Recombinants are indicated in parentheses below each SSR marker. (B) Gene structure of *ZmEHD1*. Black boxes indicate exons, and lines between black boxes represent introns. The positions of mutations were marked. The mutations that lead to the amino acid change between the *ehd1* mutant and the WT are indicated with red letters. (C) Schematic representation of the predicted structure of ZmEHD1. The regions encoding the potential protein domains are shown. The positions of mutations on coil-coil domain were marked by asterisks. (D) Real-time RT-PCR detection of *ZmEHD1* gene transcripts in endosperms of the WT maize and the *ehd1* mutant at 15, 20, 25, and 30 DAP. Quantifications were normalized to the expression of 18S rRNA. Values are means and standard errors (n=3). ** indicates a significant difference (P < 0.01) between the WT and the *ehd1* mutant.

The genomic DNA sequence of GRMZM2G052740 spanned ~4.4-kb and generated a transcript that included 16 exons (Figure 3B). The corresponding 2,209-bp cDNA sequence encoded a polypeptide of 547 amino acids with a molecular mass of ~61 kD (Supplemental Figure 1). BLASTP analysis in GenBank indicated that GRMZM2G052740 is closed related to the *Arabidopsis* EHD-containing proteins, AtEHD1 and AtEHD2 (Supplemental Figure 1). The ZmEHD1 protein shares the typical structure of the EHD family; it has an EH domain with two calcium-binding EF-hands (15-39 aa and 49-84 aa), a P-loop (GQYSTGKT), a dynamin-type guanine nucleotide-binding domain, and a coil-coil domain (Figure 3C).

According to the publicly available maize microarray database (www.qteller.com, Waters et al., 2011), *ZmEHD1* is expressed in various tissues and its mRNA abundance is high in immature cobs (V18), embryos, and endosperms at 12 and 14 DAP. To verify the microarray data, we performed quantitative real-time RT-PCR using RNA isolated from various maize tissues, including roots, leaves, stems, endosperms, and embryos, and the results confirmed the public microarray data (Supplemental Figure 2). We also isolated RNA from the endosperms of the *ehd1* mutant and the WT at 15, 20, 25, and 35 DAP to assess *ZmEHD1* expression levels. The transcripts of *ZmEHD1* were down-regulated during endosperm development in both the *ehd1* mutant and the WT (Figure 3D). Unexpectedly, *ZmEHD1* mRNA abundance at all sampling times in the *ehd1* mutant was higher than in the WT (Figure 3D), indicating that the mutated *ZmEHD1* might be only partly functional.

### Validating that *ZmEHD1* is the causative locus by *ZmEHD1* loss-of-function mutants

To determine whether *ZmEHD1* is responsible for the EHD1 locus, we used the CRISPR/Cas9 system to generate *ZmEHD1* loss-of-function mutant (KO). The given gene sequencing in *ZmEHD1* loss-of-function mutants revealed a deletion of 775-bp or 785-bp fragment at the coding sequence, which resulted in a frameshift (Supplemental Figure 3). *ZmEHD1* loss-of-function also led to decreased germination rate as well as retarded vegetative development (Figure 4A and 4B).

**Figure 4.**
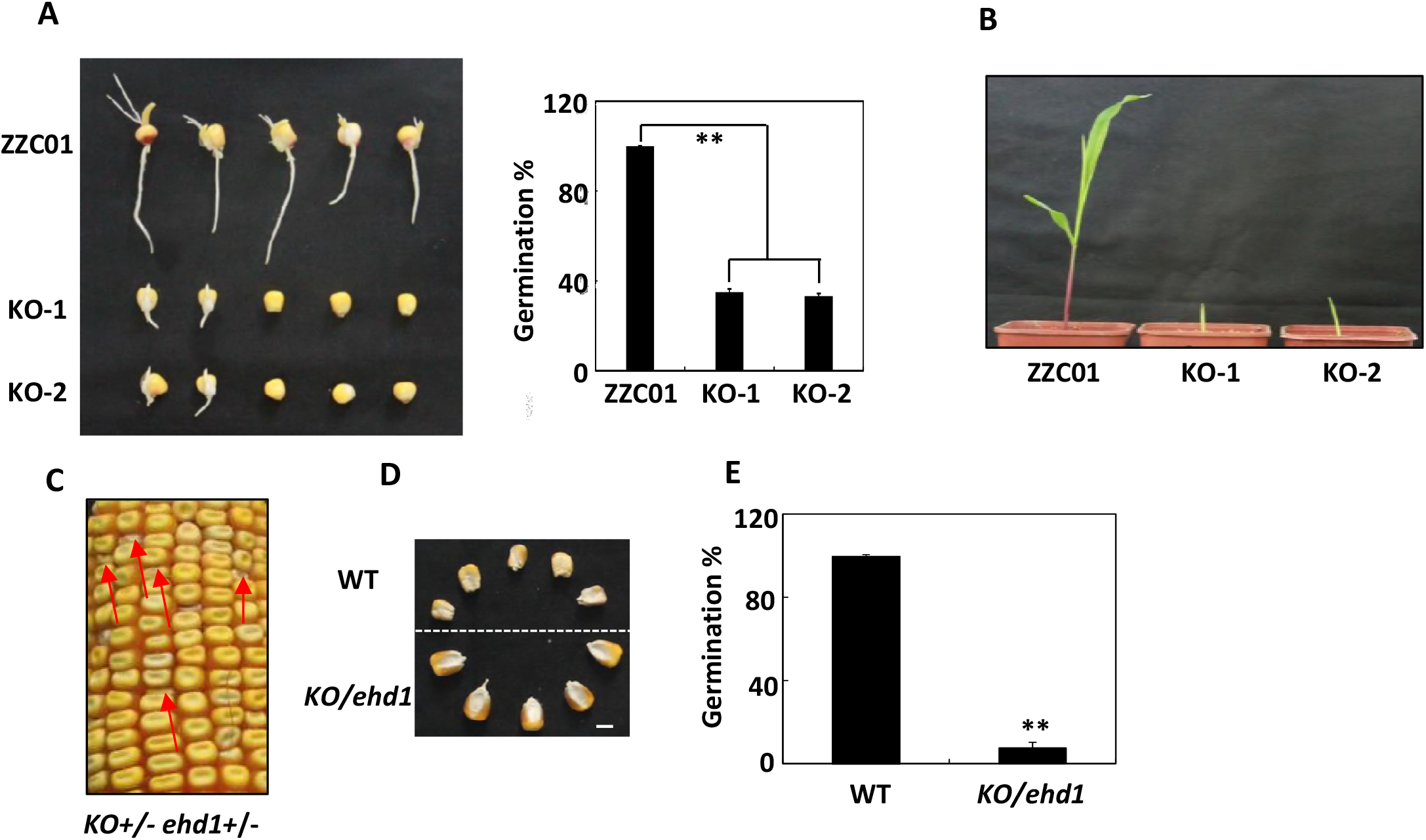
Transgenic validation of *ZmEHD1*. (A) Germination of inbred line ZZC01 (WT) and *ZmEHD1* knock-out mutants (KO). Values are means and standard errors of approximately 100 seeds from three independent experiments. An LSD test was used to assess differences between WT and *ZmEHD1* knock-out mutant. **, P < 0.01 (t-test), significant difference between WT and KO mutants. (B) Phenotypes of WT and KO mutants (n=20). Representative photograph is shown. (C) Heterozygous KO(+/−) × *ehd1* (+/−) were used in an allelism test. The red arrows indicated the *ehd1* kernels. (D) The mature kernels of WT and *KO/ehd1* mutant randomly selected from ears of KO(+/−) × ehd1 (+/−). Scale bars = 0.5 cm. (E) Germination of WT and *KO/ehd1* mutant. Values are means and standard errors of approximately 140 seeds from three independent experiments. An LSD test was used to assess differences between WT and *KO/ehd1* mutant. **, P < 0.01 (t-test), significant difference between WT and *KO/ehd1* mutant.

An allelism test was performed by crossing KO-1 F_1_ (KO+/−) and *ehd1* heterozygotes (ehd1+/−). The shrunken kernel phenotype (*KO/ehd1*) and the WT phenotype (*EHD1/EHD1*; *EHD1/KO*; *EHD1/ehd1*) in the hybrid F_1_ ears displayed a 1:3 segregation ratio (Figure 4C, 4D and Supplemental Table 4), indicating that *KO* cannot complement *ehd1*. Meanwhile, the germination rate of the *KO/ehd1* mutant was much lower than that of the WT (Figure 4E). These results indicate that *ZmEHD1* is the locus affected by *ehd1*.

### The *ehd1* and KO mutant are defective in endocytosis

EHD has usually been identified among proteins involved in endocytosis, vesicle transport, and signal transduction (Lin et al., 2001). To test whether the ZmEHD1 protein is involved in CME in maize, we examined the uptake of N-(3-triethylammonium-propyl)-4-(4-diethylaminophenylhexatrienyl) pyridinium dibromide (FM4-64), a commonly used endocytosis tracer, in the *ehd1* mutant and the WT. Because maize seedlings are relatively large, we could not place the whole plant on the microscope platform as was previously done with 3-day-old *Arabidopsis* plants (Fan et al., 2013). Instead, we detached the similar parts of roots from the *ehd1* mutant and the WT seedlings that had been treated with 5 μM FM4-64 for 10 min to monitor the FM4-64 endocytosis. At 30 minutes after labeling, FM4-64-labelled fluorescent puncta were detected in 28% of the WT root cells (21 of 75 cells), while no FM4-64-labelled fluorescent puncta were detected in *ehd1* root cells (0 of 90 cells) (Figure 5A). Although FM4-64-labelled fluorescent puncta could be detected in root cells of both the WT and the *ehd1* mutant at 60 minutes after labeling, FM4-64-labelled fluorescent puncta were detected in only 19% of the *ehd1* root cells (22 of 117 cells) but in 45% of the WT root cells (45 of 101 cells) (Figure 5A). The effects of ZmEHD1 on endocytosis were further confirmed by the delayed internalization of FM4-64 in *ZmEHD1* knock-out mutants (Figure 5B).

**Figure 5.**
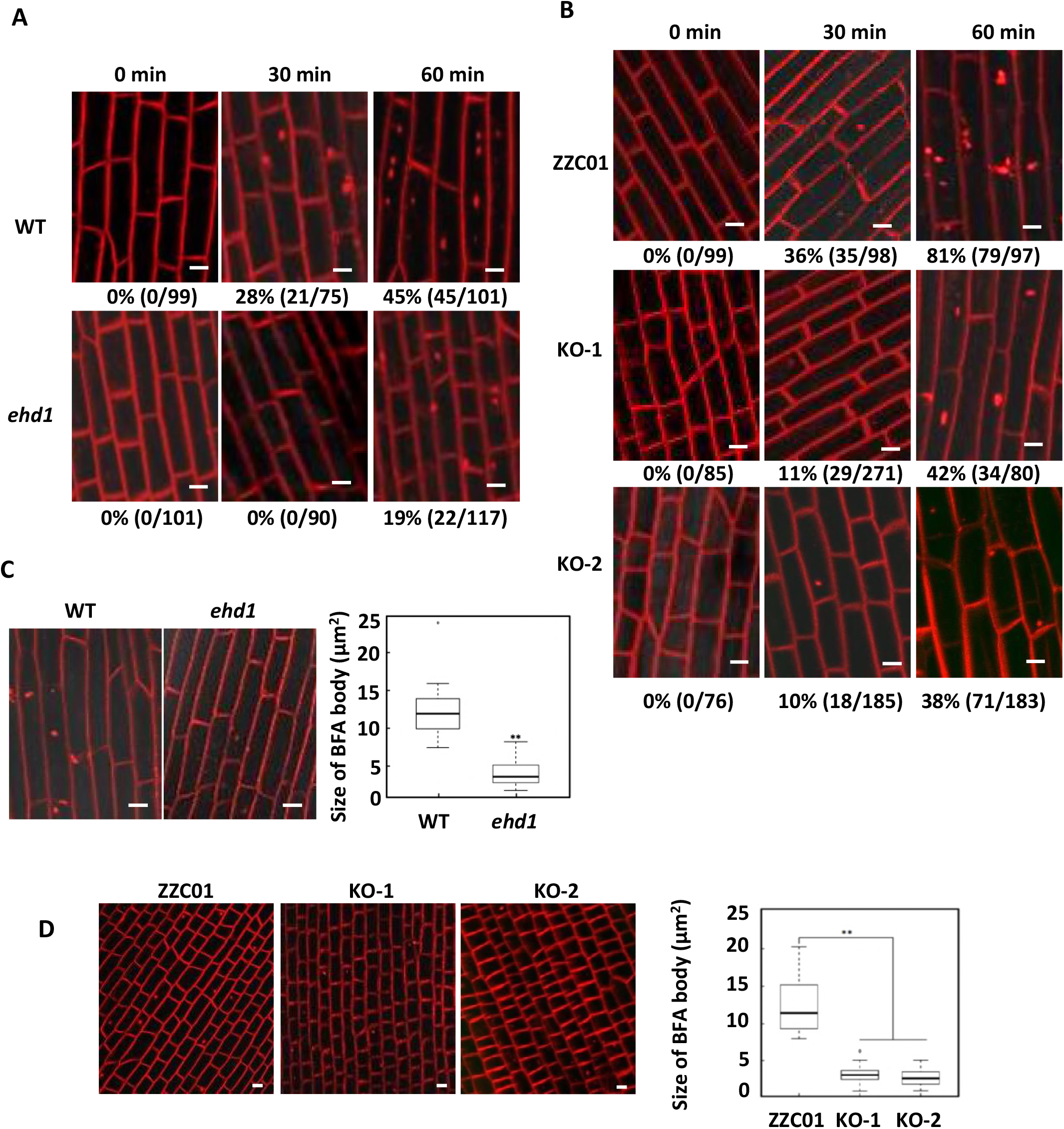
Endocytosis of FM4-64 is reduced in the *ehd1* and in *ZmEHD1* knock-out mutants. Three dimensional reconstructions of z-stacks (60 μM with 2.4-μM steps) were obtained in the *ehd1* (A) and in *ZmEHD1* knock-out mutant (B). FM-64 labeled BFA bodies in the *ehd1* mutant (C) and in *ZmEHD1* knock-out mutants (D). Representative photographs at indicated durations are shown. Scale bars = 10 μm. Numbers below the photographs indicate the rate of FM4-64-labelled fluorescent puncta (% and number of FM4-64-labelled fluorescent puncta/total cell ** number). indicates a significant difference from the WT at P < 0.01 according to a *t*-test.

To further investigate whether ZmEHD1 was involved in the vesicle traffic pathway, we inhibited endocytic recycling of FM-64 using the fungal toxin brefeldin A (BFA). The accumulation of FM-64 in BFA bodies was clearly observed in the WT when treated with BFA (Figure 5C and 5D). In contrast, the aggregation of FM-64 in BFA bodies remarkably decreased in the *ehd1* mutant and *ZmEHD1* knock-out mutants (Figure 5C and 5D). Compared to the WT, the sizes of FM-64 labeled BFA bodies in the *ehd1* mutant and *ZmEHD1* knock-out mutants decreased by ~67% and ~76%, respectively (Fig. 5C and 5D). These results suggested that ZmEHD1 is important in endocytic transport.

### Subcellular localization of ZmEHD1 and ZmEHD1_mut_

To determine the subcellular localization of ZmEHD1 protein, yellow fluorescent protein (YFP) fused N-terminally to ZmEHD1 was transiently expressed in tobacco (*N. benthamiana*) leaves under the control of the *CaMV 35S* promoter. In tobacco epidermal cells expressing the YFP-ZmEHD1 fusion protein, the YFP signal was mainly detected in the PM (Supplemental Figure 4A). The localization was confirmed by staining lipid membranes with the red fluorescent probe, FM4-64 (Supplemental Figure 4A). We also cloned the *ZmEHD1* CDS from the *ehd1* mutant and examined the subcellular localization of ZmEHD1_mut_ protein. YFP-ZmEHD1_mut_ signals were aggregated and colocalized with FM4-64 signals in tobacco epidermal cells (Supplemental Figure 4B). This phenomenon was consistently observed in different biological replicates. Thus, we deduced that the mutation in ZmEHD1 affected the subcellular localization of ZmEHD1.

### ZmEHD1 interacts with the ZmAP2 σ subunit in yeast and plant

In its drastically reduced fertility and endocytosis, the *ehd1* mutant resembles *Arabidopsis ap2 σ* mutants, in which CME and multiple stages of plant development are impaired (Fan et al., 2013). We hypothesized that ZmEHD1 is involved in CME by virtue of its interaction with the *ZmAP2 σ* subunit. To test this hypothesis, we cloned the cDNA sequence of the maize AP2 σ subunit (GRMZM2G052713) and introduced it into *Arabidopsis ap2 σ* loss-of-function mutant (Fan et al., 2013). *Arabidopsis ap2 σ* homozygous for the T-DNA insertion was identified by PCR (Supplemental Figure 5A). RT-PCR showed that *ZmAP2 σ* subunit was expressed in the *Arabidopsis ap2 σ* mutant (Supplemental Figure 5B). The transformation of the *Arabidopsisap2 σ* mutant with *ZmAP2 σ* subunit rescued the developmental defects, such as reduced leaf size and fertility (Supplemental Figure 5C). These results suggested that *ZmAP2 σ* subunit has similar functions as *AtAP2 σ* subunit in CME.

The potential interaction between ZmEHD1 and the ZmAP2 σ subunit was first evaluated by a biomolecular fluorescence complementation (BiFC) assay in tobacco leaves and maize mesophyll protoplasts. The ZmEHD1, ZmEHD1_mut_, and ZmAP2 σ subunit constructs were introduced into *N. benthamiana* mesophyll cells or maize mesophyll protoplasts by *A. tumefaciens*-mediated infiltration. The vitality of the *N. benthamiana* mesophyll cells was determined by propidium iodide. Reconstituted YFP signals were observed in the PM when both nYFP-ZmAP2 σ subunit and cYFP-ZmEHD1 constructs were co-expressed (Figure 6A and Supplemental Figure 6). Reconstituted YFP signals were also observed when the nYFP-ZmAP2 σ subunit and cYFP-ZmEHD1_mut_ constructs were co-expressed (Figure 6A and Supplemental Figure 6). Consistent with the subcellular localization of ZmEHD1mut, the reconstituted YFP signals were diffused throughout the cells when the nYFP-ZmAP2 σ subunit and cYFP-ZmEHD1_mut_ constructs were co-expressed (Figure 6A and Supplemental Figure 6). This was further verified by the localization of ZmAP2σ subunit in WT maize and *ehd1* mutant (Supplemental Figure 7).

**Figure 6.**
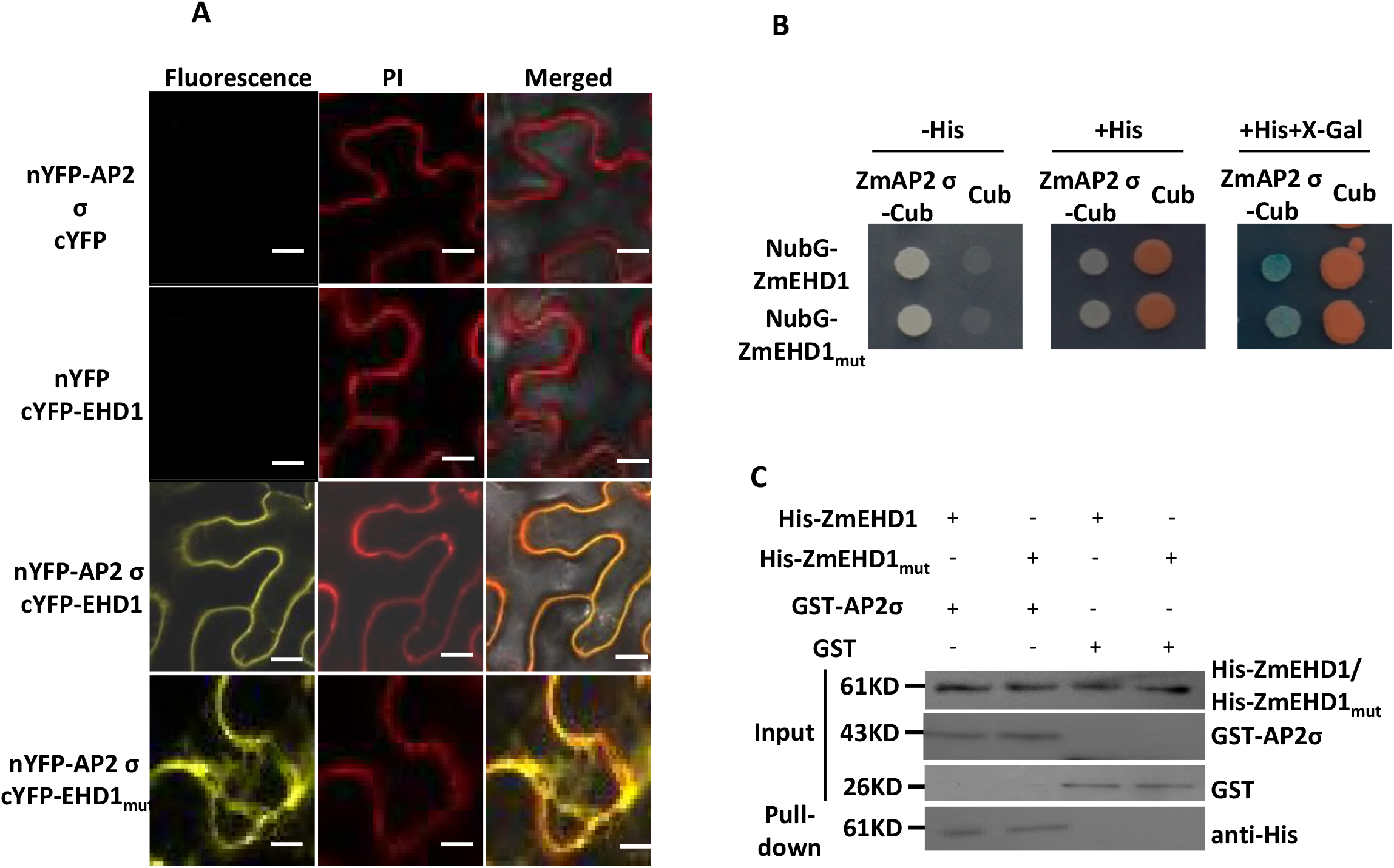
ZmEHD1 directly interacts with the ZmAP2 σ subunit in plants and yeast. (A) Interaction between ZmEHD1 and the ZmAP2 σ subunit in the leaves of *N. benthamiana*as observed by BiFC. The photographs were taken in the dark field for yellow fluorescence. Propidium iodide (PI) was used to determine the vitality of the cells. Representative photographs are shown. Scale bars = 10 μm. (B) Interaction between ZmEHD1 and the ZmAP2 σ subunit as indicated by split-ubiquitin yeast two-hybrid assays. The ZmAP2 σ subunit was used as the fused bait protein (AP2 σ-Cub), and ZmEHD1 was used as the fused prey protein (NubG-ZmEHD1). The presence or absence of His or X-Gal is indicated. (C) Pull-down assays for the interaction between ZmEHD1 and the ZmAP2 σ subunit. The ZmEHD1/ ZmEHD1_mut_ in the pull-downed fraction was detected by immunoblot using anti-His antibody.

We also used a split-ubiquitin membrane-based yeast two-hybrid system to verify the interaction between the ZmAP2 σ subunit and ZmEHD1 as well as ZmEHD1_mut_. Both ZmEDH1 and ZmEHD1_mut_ interacted directly with ZmAP2 σ subunit in yeast. The interaction resulted in survival in a medium lacking His and β-galactosidase activity (Figure 6B).

Pull-down experiments using purified His-ZmEHD1/His-ZmEHD1mut to isolate GST-ZmAP2 σ subunit also confirmed the physical interaction between ZmEHD1/ZmEHD1_mut_ and ZmAP2 σ subunit. His-ZmAP2 σ subunit was clearly detected in the pull-down fraction (Figure 6C). These results strongly suggested that ZmEHD1 directly interacts with ZmAP2 σ subunit at the plasma membranes in maize.

### Transcriptome profiling of the *ehd1* mutant

To gain insight into the molecular events involved in the *ZmEHD1* -mediated signaling pathway, we compared the whole-transcriptome profiles of endosperms of the *ehd1* mutant and the WT at 15 DAP using RNA-seq. Each sample was represented by two biological replicates, and the libraries were sequenced by Illumina high-throughput sequencing technology. These four RNA libraries yielded more than 0.5 billion raw reads, and approximately 97% of the raw reads remained after adaptor polluted reads, Ns reads, and low quantity reads were trimmed. Of the remaining reads, more than 0.12 billion could be perfectly mapped to maize B73 RefGen_V3.27 (ftp://ftp.ensemblgenomes.org/pub/plants/release-27/fasta/zea_mays). Sequences that could not be mapped to the maize genome were discarded, and only those perfectly mapped were analyzed further. The abundance of each gene was expressed by reads per kilo base million mapped reads (RPKM) (Wagner et al., 2012). We calculated the Pearson’s correlation coefficients (*R* value) of the two biological replicates for each treatment to investigate the variability between the replicates. The *R* value of both comparisons exceeded 0.99 (Supplemental Figure 8), indicating a high correlation between biological replicates.

Based on the criteria that the Log_2_ fold-change ratio was ≥ 1 and that the adjusted P value was ≤ 0.05, 4,760 genes, including *ZmEHD1*, were identified as differentially expressed genes (DEGs) by DEGseq software (Wang et al., 2010). Among the DEGs, 2,208 genes were up-regulated and 2,552 genes were down-regulated in the *ehd1* mutant relative to the WT (Supplemental Table 5). The results of RNA-seq were confirmed by quantitative real-time RT-PCR. In agreement with our RNA-seq data, the expression levels of randomly selected GRMZM2G088273, GRMZM2G346897, GRMZM2G067929, and GRMZM2G418119 were lower in the *ehd1* mutant than in the WT (Supplemental Figure 9A). As expected, GRMZM2G156877 and GRMZM2G420988 were expressed at higher levels in the *ehd1* mutant than in WT (Supplemental Figure 9A), demonstrating the reliability of our RNA-seq data. We then performed Gene Ontology (GO; http://bioinfo.cau.edu.cn/agriGO/) analysis to determine the molecular events related to the DEGs. GO analysis indicated that the 4,760 DEGs were highly enriched for biological processed involved in response to stress (GO:0006950, P = 1.2e^-12^), integral to membrane (GO:0016021, P = 1e^-5^), intracellular membrane-bounded organelle (GO:0043231, P = 0.0002), vacuole (GO:0005773, P = 0.0007), starch metabolic processes (GO:0005982, P = 1.6e^-5^), and carbohydrate metabolic processes (GO:0005975, P = 4.16e^-5^).

Interestingly, some of the DEGs have known or presumed functions associated with auxin-mediated signaling pathway (GO:0009734, P = 1.2e^-25^) and response to auxin stimulus (GO:0009733, P = 2.1e^-25^), including AUXIN RESPONSE FACTOR (ARF), AUX/IAA transcription factor, small auxin up-regulated RNA (SAUR), indole-3-acetaldehyde oxidase, auxin transporters, and efflux carrier (Table 1). We randomly chose four of the DEGs listed in Table 1 (GRMZM2G082943, GRMZM2G365188, GRMZM5G809195, and GRMZM2G019799) and validated their difference in expression levels in the *ehd1* mutant vs. the WT by quantitative real-time RT-PCR (Supplemental Figure 9B). These results indicated that *ZmEHD1* might affect auxin homeostasis in maize.

### Auxin distribution and ZmPIN1a-YFP localization in the *ehd1* mutant

To test the hypothesis that *ZmEHD1* might affect auxin homeostasis in maize, we first explored the gravitropic response by examining mesocotyl-coleoptiles. We placed the upright mesocotyl-coleoptiles in the horizontal direction. In contrast to less than 4 h for the recovered vertical growth of WT, it took about 5 h for the *ehd1* mutant and *ZmEHD1* knock-out mutants to recover vertical growth under the same condition (Figure 7A). The free IAA content in kernels of the *ehd1* mutant and the WT at 15 DAP were determined. The free IAA content in the kernels of the *ehd1* mutant was 2,558 pg mg^-1^ FW, which was ~30% lower than that in the WT (Figure 7B).

**Figure 7.**
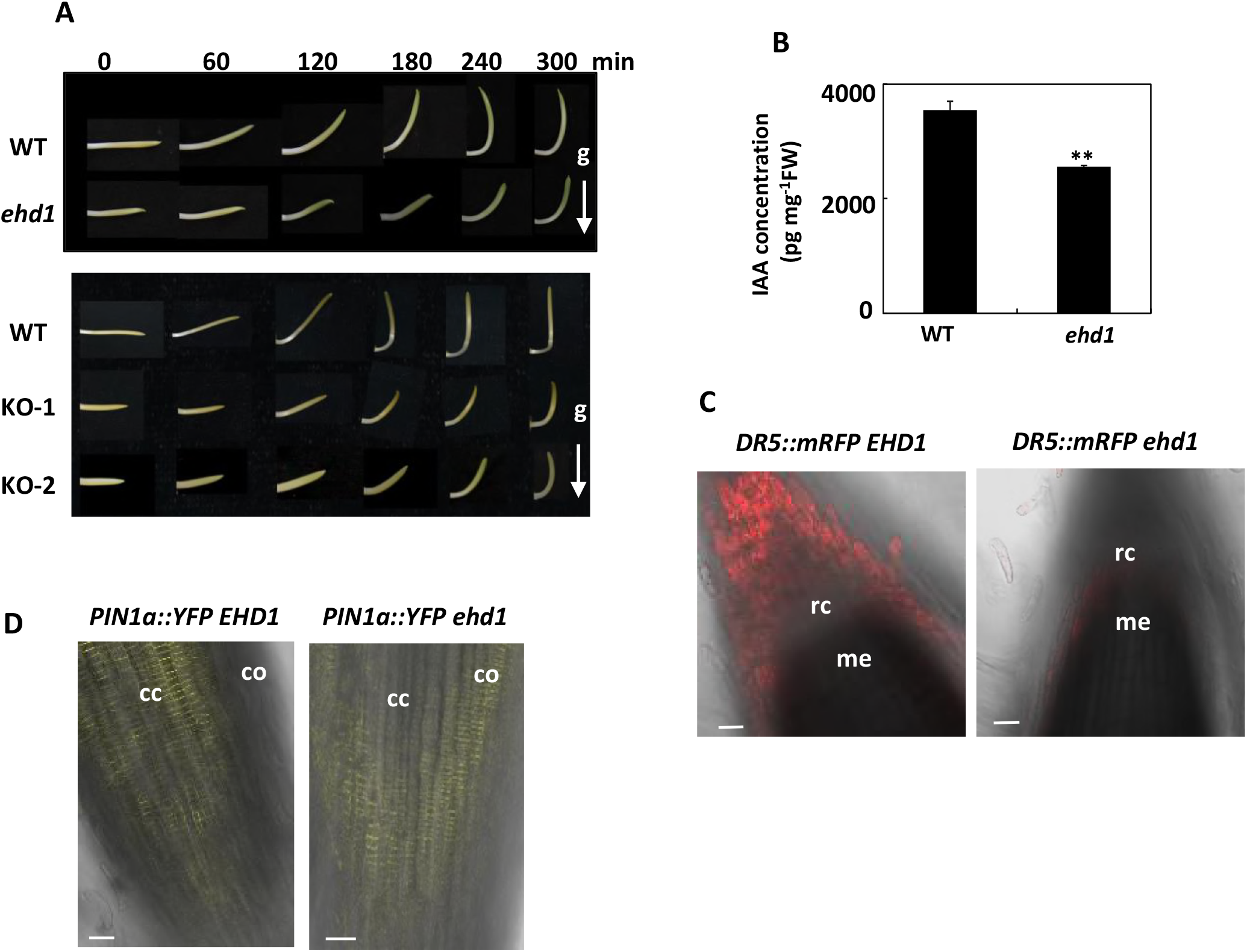
Auxin distribution and ZmPIN1a-YFP localization were altered in the *ehd1* mutant. (A) The response of horizontally placed mesocotyl-coleoptiles to gravity is delayed in *ehd1* mutant (n=15). (B) Free IAA contents in WT maize and *ehd1* mutant kernels at 15 DAP. Error bars represent standard errors (n=4). ** indicates a significant difference from the WT at P < 0.01 according to a t-test. (C) The fluorescence of ZmDR5::mRFP in WT and *ehd1* mutant. Scale bars = 50 μm. (D) The fluorescence of ZmPIN1a::YFP in WT and *ehd1* mutant. Scale bars = 50 μm. cc, central cylinder; me, root meristem; rc, root cap; co, cortex.

To further demonstrate that ZmEHD1 regulates auxin homeostasis in maize, we first crossed the auxin-responsive ZmDR5::RFP reporter maize with *ehd1* mutant to examine the auxin distribution in the root tips of *ehd1* mutant. The RFP signals were significantly reduced in the root caps in *ehd1* mutant (Figure 7C). We also crossed the ZmPIN1a::YFP line with *ehd1* mutant. The YFP signals in the roots of *ehd1* mutant showed more diffuse localization than that in *PIN1a::YFP EHD1* line, and could be detected in root cortex (Figure 7D). These results suggested that auxin homeostasis was altered in *ehd1* mutant.

### 1-NAA application rescues the fertility of the *ehd1* mutant

To further verify that the rather low fertility of the *ehd1* mutant and *ZmEHD1* knock-out mutant was caused by auxin homeostasis, we attempted to rescue the phenotypic defects of *ehd1* mutant and *ZmEHD1* knock-out mutants by exogenous application of the active auxin compound, 1-naphthaleneacetic acid (1-NAA). The seeds of the *ehd1* mutant and *ZmEHD1* knock-out mutants were rinsed in water or in four increasing concentrations of 1-NAA for 12 h, and then germinated in the paper-culture system. Little or no phenotypic rescue of the *ehd1* mutant was observed when 30 mg L^-1^ 1-NAA or water alone was applied (Figure 8A). However, the germination rate of *ehd1* mutant and *ZmEHD1* knock-out mutant was significantly increased when treated with low concentrations of 1-NAA (Figure 8A). Also, when treated with low concentrations of 1-NAA, the *ehd1* mutant and *ZmEHD1* knock-out mutant survived after its first two leaves had completely expanded (Figure 8B). In contrast to IAA, exogenous GA3 had no effect on the germination of the *ehd1* mutant and *ZmEHD1* knock-out mutant (Supplemental Figure 10). Overall, these results demonstrated that the phenotypic defects of the *ehd1* mutant could be rescued by exogenous application of 1-NAA.

**Figure 8.**
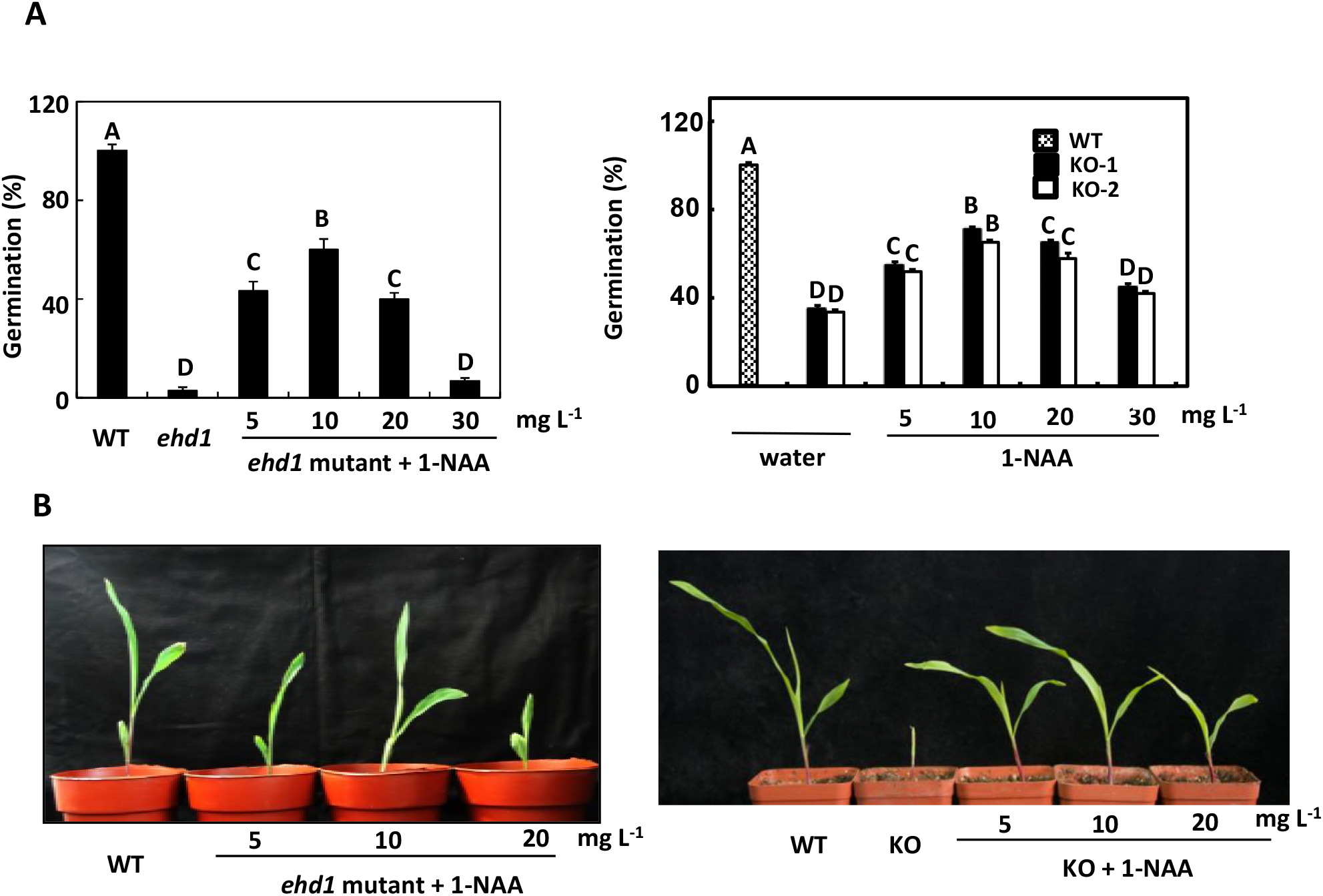
Rescue of the *ehd1* phenotype by exogenous 1-NAA. (A) Germination of the *ehd1* mutant *ZmEHD1* knock-out mutants (KO) after treatment with water alone or with different concentrations of 1-NAA. Values are means and standard errors of approximately 100 seeds from three independent experiments. An LSD test was used to assess differences between treatments. Means with the same letter are not significantly different at P < 0.01. (B) Phenotypes of *ehd1* and KO seedlings treated with different concentrations of 1-NAA.

## Discussion

In the present research, we characterized the *ehd1* mutant, which is impaired in kernel development and vegetative growth, and used positional cloning and transgenic validation to verify that the *ehd1* gene encodes an EHD protein. The results of an FM4-64 uptake experiment, split-ubiquitin membrane-based yeast two-hybrid system, BiFC and pull-down assay indicated that ZmEHD1 is involved in CME through its interaction with the ZmAP2 σ subunit. Additionally, the transcriptome profiling of the ehd1 mutant, the auxin distribution and ZmPIN1a-YFP localization in the *ehd1* mutant, and the rescue of the mutant phenotypes following the exogenous application of 1-NAA revealed that the ZmEHD1-mediated endocytosis mainly affects auxin homeostasis in maize.

In *Arabidopsis*, AP2 σ subunit is primarily recruited from the cytoplasm to the PM to initiate clathrin-coated endocytic vesicle formation (Fan et al., 2013; Fan et al., 2015). The developmental defects of *ap2 σ* loss-of-function *Arabidopsis*, such as reduced leaf size and fertility, could be rescued by *ZmAP2 σ* cDNA, suggesting that *ZmAP2 σ* subunit has similar functions as *AtAP2 σ* subunit in CME. Microarray data revealed that the AP2 σ subunit is ubiquitously expressed in various tissues throughout maize development (Sekhon et al., 2011), which is consistent with the expression pattern of *ZmEHD1*. Based on the direct interaction between the ZmAP2σ subunit and ZmEHD1 at the PM as demonstrated by BIFC, pull-down assay and split-ubiquitin membrane-based yeast two-hybrid system in the current study, we suspected that ZmEHD1 contributes to CME by interacting with the ZmAP2 σ subunit.

The naturally occurring *ehd1* mutant had amino acid substitutions in three positions. Of these, the V/A substitution was in a linker region. The other two mutations were in the coil-coil domain of ZmEHD1. The coil-coil domain was responsible for protein-protein interactions (Bar et al., 2009; Sharma et al., 2008).

Here, we showed that the subcellular location of ZmEHD1_mut_ was obviously different from that of ZmEHD1. Though ZmEHD1_mut_ could directly interact with ZmAP2σ subunit, the PM was not the main location for interaction between ZmEHD1_mut_ and the ZmAP2σ subunit. The impaired clathrin-coated endocytic vesicle formation and/or the recycling of the ZmAP2 σ subunit from the endosome to the PM should be the major contributor to phenotypes observed in the *ehd1* mutant.

*Arabidopsis ARF2* controls seed size by repressing cell proliferation (Schruff et al., 2006); in rice, activation of *BIG GRAIN1* (BG1) significantly improves grain size by regulating auxin transport (Liu et al., 2015); *TGW6*, an IAA-glucose hydrolase, negatively regulates endosperm development of rice (Ishimaru et al., 2013). Analyses of a seed-specific viable maize mutant, defective endosperm 18 (*de18)*, demonstrated that the *ZmYUC1* gene (which is critical for IAA biosynthesis) is essential for normal endosperm development in maize (Bernardi et al., 2012). Thus, auxin biosynthesis, transport and signaling might coordinate to regulate vegetative and reproductive development in maize (Gallavotti et al., 2008; Li and Li, 2016). The current results with maize show that *ARFs*, indole-3-acetaldehyde oxidase, auxin transporters, and efflux carrier are differentially expressed in the *ehd1* mutant vs. the WT; that the concentration of free IAA is lower in the *ehd1* mutant than in the WT; that the auxin distribution and ZmPIN1a-YFP localization were altered in the *ehd1* mutant; and that exogenous application of 1-NAA rescues the phenotypes of the *ehd1* mutant. These results suggest that the growth defects of the *ehd1* mutant result from the loss of auxin homeostasis.

## Materials and methods

### Plant materials

The ZmPIN1a::YFP and ZmDR5::mRFP lines were kindly provided by Lixing Yuan (China Agricultural University). The maize mutant ehd1 was isolated by screening for natural mutants defective in grain filling. To construct mapping population, the mutant was crossed with inbred line Xun9058 in the winter of 2011 in Sanya, Hainan Province. Xun9058 is the male parent of the elite hybrid Xundan20 and has been widely used in the breeding of hybrid maize in China. F_2_ individuals were obtained by selfing the F_1_ plants in the summer of 2012 in Zhengzhou, Henan Province. The site (113°42’E, 34°48’N) is located in central China and has an average annual temperature of 14.3 °C and an average annual rainfall of 640.9 mm. A F_2_ population with 165 individuals was used to map the preliminary location of *ZmEHD1* gene. Normal kernels from F_3_ segregation ears were used for fine mapping of the candidate gene. The F_3_ population contains ~53,000 individuals. A small piece of each F_3_ kernel was chipped and genotyped before planting. At harvest stage, the ear phenotype was investigated to verify the kernel segregation of each individual. Two markers, bnlg589 and RM1, were closely linked to the locus to determine recombined individuals.

From F_2_ generation, the normal kernels from segregation ears were analyzed by closely linked marker with *ZmEHD1* gene and the recombined individuals were selfed. This process was continued for five generations to construct nearly isogenic lines (NILs). The NILs were used as WT.

### Molecular markers

Bulked segregant analysis (BSA) was used to detect the genetic linkage of the *ZmEHD1* gene (Michelmore et al., 1991) and 1082 SSR markers from the maize genome database (www.maizegdb.org) were tested for polymorphism in the two parents and the two bulks. To develop new markers for fine mapping, the sequence of the B73 reference genome between markers bnlg589 and umc2289 on chromosome 4 was downloaded from MaizeGDB (http://www.maizegdb.org/). SSR-Hunter software was used find simple sequence repeats (SSRs). After BLAST was performed (http://blast.ncbi.nlm.nih.gov/Blast.cgi), SSRs with single copy were developed to new markers with PRIMER 5.0. The newly designed SSRs were tested by polyacrylamide gel electrophoresis (PAGE) for polymorphism in the *ehd1* mutant, Xun9058, pooled samples of normal kernels (*EHD1/EHD1* and *EHD1/ehdl*) or shrunken kernels (*ehdl/ehdl*). SSRs with polymorphism were used for subsequent fine mapping. The sequences of the primers used for mapping are listed in Supplemental Table 3.

### Transgenic functional validation

The CRISPR/Cas9-mediated *ZmEHD1* editing was performed as described by Xing et al. (2014) with some modifications. In brief, two gRNAs that direct target sequences located at nucleotides 16 and 791 of *ZmEHD1* were produced using primers listed in Supplemental Table 3. Thereafter, the fragments were cloned into the pBUE411 vector using the *Bsa*I restriction site by T4 Ligase reaction. The plasmid contained the *Streptomyces* hygroscopicus phosphinothricin acetyltransferase gene (bar) under the control of a *CaMV 35S* promoter and was electroporated into *A. tumefaciens* EHA105. Immature embryos of maize inbred line ZZC01 were transformed by co-cultivation with EHA105 at Life Science and Technology Center of China National Seed Group Co., LTD. Transformants were selected with gradually increasing concentrations of Bialaphos. T_2_ homozygous lines were sequenced to ensure that *ZmEHD1* was knocked out.

For *Arabidopsis* transformation, the cDNAs of *ZmAP2 σ* subunit without 3’ UTR were amplified by PCR. The corresponding products were introduced into the pENTR^TM^/D-TOPO vector (Invitrogen) and cloned into pMDC32 by LR reactions (Invitrogen). The plasmid was electroporated into *A. tumefaciens* GV3101 and was transformed into *Arabidopsis* by the floral dip method (Clough and Bent, 1998). Transgenic plants were selected with the use of 35 μg ml^-1^ hygromycin. The sequences of the primer pairs used in the experiments are listed in Supplemental Table 3.

### Expression analysis of *ZmEHD1*

Total RNA was extracted from the WT and the *ehd1* mutant with the RNeasy plant mini kit. The RNA was used for the synthesis of first-strand cDNA with SuperScript III first-strand synthesis supermix (Invitrogen). Quantitative real-time RT-PCR was carried out in an ABI 7500 system using the SYBR Premix Ex TaqTM (perfect real time) kit (TaKaRaBiomedicals), with 18S rRNA and *Actin* as controls. Primers designed to detect the transcription level of *ZmEHD1* were listed in Supplemental Table 3. The relative expression level was calculated using the comparative Ct method. Each experiment was replicated at least three times.

### Subcellular localization of ZmEHD1

The full-length *ZmEHD1* and *ZmEHD1_mut_* coding region was amplified from the WT and the *ehd1* mutant. To generate the *ZmEHD1* expression plasmids, the 1644-bp fragments were cloned into the pCPB vector, which was fused to an YFP protein in the N terminus using the *Bam*HI restriction site by In-Fusion reaction. The plasmid was electroporated into A. *tumefaciens* GV3101 and was transiently expressed in tobacco epidermal cells as described previously (Li et al., 2008). To stain the PM of epidermal cells, 5 μM FM4-64 was infiltrated into tobacco leaves for 3 h before observation. YFP/FM-64 images were collected with a Zeiss LSM700 confocal microscope.

### Split-ubiquitin yeast two-hybrid assay

The split-ubiquitin two-hybrid system was used to detect the interactions between membrane proteins. The assay was carried out according to the instructions provided with the DUALmembrane Kit (Dualsystems Biotech). The full-length coding regions of the *ZmAP2* σ subunit, *ZmEHD1* and *ZmEHD1_mut_* were amplified and cloned into the vectors pST3-STE and pPR3-N NubG using the *SfiI* restriction site by In-Fusion reaction. Vectors were co-transformed into yeast strain NMY51. The interactions were assessed by the growth of yeast colonies on synthetic minimal medium containing 7.5 mM 3-aminnotriazole without Leu, Trp, His and Ade and also by chloroform overlay β-galactosidase plate assay (Duttweiler, 1996).

### Bimolecular fluorescence complementation (BiFC) assays

The coding sequences of the *ZmAP2* σ subunit, *ZmEHD1* and *ZmEHD1_mut_* were amplified using specific primers listed in Table S1, and were cloned into the pSPYNE-35S or pSPYCE-35S binary vectors using the *BamHI* restriction site. The various combinations of *ZmAP2* σ subunit, *ZmEHD1*, and *ZmEHD1_mut_* expression vectors were transiently expressed in 3-week-old *N. benthamiana* leaves by *Agrobacterium-mediated* infiltration (strain EHA105). YFP images were obtained 3 days after infiltration with a Zeiss LSM700 confocal microscope.

### Pull-down analysis

The full-length coding sequences of *ZmEHD1, ZmEHD1_mut_* and *ZmAP2* σ subunit were individually subcloned into the pET-28a(+) and pGEX-4T-1 vectors using the *Bam*HI restriction site by In-Fusion reaction. The resulting constructs were verified by sequencing. His-ZmEHD1, His-ZmEHD1_mut_, GST and GST-ZmAP2 σ subunit were expressed in *Escherichia coli* BL21.

For the *in vivo* pull-down analysis, 10 μg of purified His-ZmEHD1, His-ZmEHD1mut were incubated with GST-ZmAP2 σ subunit or GST bound to glutathione-Sepharose™ 4B (GE Healthcare) for 4 h at 4 °C on a rotary shaker. Precipitated beads were washed six times with washing buffer. Washed beads were boiled with 50 μL of 1 × SDS sample buffer for 5 min and subjected to SDS-PAGE and immunoblot analysis.

### FM4-64 internalization assay

FM4-64 internalization assays were carried out to evaluate the rate of endocytosis as described by Fan et al. (2013) with some modifications. WT and *ehd1* seedlings were incubated in half-strength Hoagland’s nutrient solution containing 5 μM FM4-64 for 10 min at room temperature. The roots were then cut and transferred to glass slides. The FM4-64 internalization was monitored at indicated durations at room temperature with a Zeiss LSM700 confocal microscope.

### RNA-seq analysis

At 15 DAP, total RNA was extracted from the endosperms of the *ehd1* mutant and the WT with TRIZOL reagent (Invitrogen), and 3 μg of total RNA was used as input material for construction of the RNA libraries. The RNA-sequencing libraries were constructed with NEBNext^®^ Ultra™ RNA Library Prep Kit for Illumina^®^ (NEB, USA). In brief, mRNA was purified from total RNA using poly-T oligo-attached magnetic beads. The enriched mRNA was then fragmented. The random hexamers were used for first-strand cDNA synthesis. After second-strand cDNA synthesis, terminal repair, and poly(A) tail and sequencing oligonucleotide adaptors ligation, the fragments were purified and subsequently amplified by PCR. The insert size was assessed with the Agilent Bioanalyzer 2100 system (Agilent Technologies, CA). Finally, the libraries of inserting cDNAs with 200-bp in size were generated and sequenced with the IlluminaHiseq 2500 platform (ANROAD, Beijing, China).

The raw reads were produced after exclusion of low quality reads and 5’ and 3’ adaptor contaminants. The unique RNAs were aligned to the maize RefGen_V3.27 (ftp://ftp.ensemblgenomes.org/pub/plants/release-27/fasta/zea_mays). Only perfectly matching sequences were considered for further analysis. The count information was used to determine normalized gene expression levels as RPKM (Wagner et al., 2012). Multiple testing with the Benjamini-Hochberg procedure for false discovery rate (FDR) was taken into account by using an adjusted P-value. Genes were considered to be differentially expressed if the Log_2_ fold-change ratio was ≥ 1 and if the adjusted P value was < 0.05 according to the DEGseq method (Wang et al., 2010).

### Free IAA analysis

At 15 DAP, kernels of the WT and the *ehd1* mutant were collected and frozen in liquid nitrogen. A 200-mg (fresh weight) sample of WT and *ehd1* kernels was finely ground in liquid nitrogen and then extracted with 1 ml of methanol containing antioxidant and ^2^H_2_-IAA (internal standard, CDN Isotopes) at 4°C for 24 h. After centrifugation, the extract was purified with an Oasis Max solid phase extract cartridge (150 mg/6 cc; Waters). IAA was quantified using UPLC-MS/MS consisting of a UPLC system (ACQUITY UPLC, Waters) and a triple quadruple tandem mass spectrometer (Quattro Premier XE, Waters) as described by Wang et al. (Wang et al., 2015). Four independent biological replicates and two technical repeats were performed for the WT and the *ehd1* mutant.

### Phytohormone treatments

For phytohormone treatments, WT and *ehd1* mutant seeds were immersed in water and the indicated concentrations of 1-NAA for 12 h or GA3 for 24 h. The seeds were then kept at 22 °C in the dark for 120 h for germination. A seed was scored as germinated if its radicle protruded through the seed coat.

## Additional information

### Funding

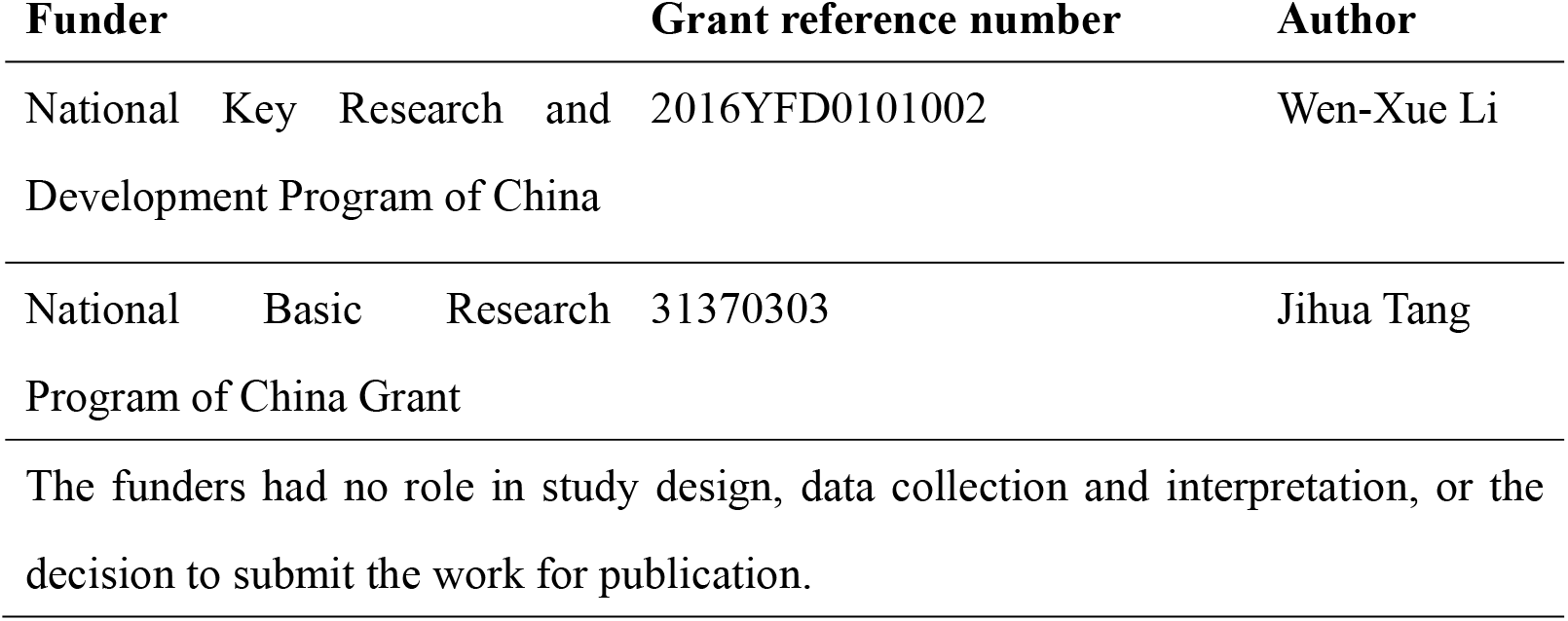

### Author contributions

J.T. and W.X.L. designed the research; Y.W., W.L., H.W., Q.D. and Z.F. performed the research; W.X.L. and J.T. analyzed the data and wrote the article.

---

**Supplemental Table 1. The ear performance of F_2_ and F_3_ individuals evaluated in the fields.**

**Supplemental Table 2. The kernel performance of F_2_ population evaluated in the fields.**

**Supplemental Table 3. Oligos used as primers in the experiment.**

**Supplemental Table 4. Allelism test between *KO+/−* and *ehd1+/−***

**Supplemental Table 5. List of genes with significant expression changes.**

**Supplemental Figure 1. The alignment with AtEHD1, AtEHD2, ZmEHD1 and ZmEHD_mut_**. The motifs are underlined.

**Supplemental Figure 2. Tissue pattern of *ZmEHD1* transcript accumulation.**

**Supplemental Figure 3. The observed target deletion of *ZmEHD1* in *ZmEHD1* loss-of-function mutants.**

**Supplemental Figure 4. The different subcellular localization of ZmEHD1 protein (A) and ZmEHD1mut protein (B). Representative photographs are shown.**

**Supplemental Figure 5. Rescue of *Arabidopsis ap2 σ* mutant by the *ZmAP2 σ* cDNA**. (A) Identification of *ap2 σ* T-DNA insertion mutant. (B) Semi-quantitative RT-PCR analysis of *ZmAP2 σ* transcript levels in the Col, *ap2 σ* mutant and *35S::ZmAP2 σ* transgenic plants. (C) Three week-old and five week-old of WT (left), *ap2 σ* mutant (middle) and *35S::ZmAP2 σ* transgenic plants.

**Supplemental Figure 6. Interaction between ZmEHD1 and the ZmAP2 σ subunit in maize mesophyll protoplasts observed by BiFC.** The photographs were taken in the dark field for yellow fluorescence. Representative photographs are shown. Scale bars = 10 μm.

**Supplemental Figure 7. The different subcellular localization of ZmAP2 σ subunit in WT maize and *ehd1* mutant.** Representative photographs are shown. Scale bars = 10 μm.

**Supplemental Figure 8. Pearson’s correlation coefficients (*R* value) of biological replicates in the WT maize and the *ehd1* mutant.**

**Supplemental Figure 9. Validation of RNA-Seq by real-time RT-PCR.** (A) Genes listed in Supplemental Table 5. (B) Genes listed in Table 1.

**Supplemental Figure 10. Germination of the *ehd1* mutant (A) and *ZmEHD1* knock-out mutant (B) treated with water alone or with different concentrations of GA3.** Values are means and standard errors of approximately 150 seeds from three independent experiments.

